# GPR182 is a lipoprotein receptor for dietary fat absorption

**DOI:** 10.1101/2025.02.03.634329

**Authors:** Zhiwei Sun, Robert J Torphy, Emily N Miller, Anza Darehshouri, Isaac Vigil, Taichi Terai, Eck Eleanor, Yi Sun, Yujie Guo, Elliott J Yee, Junyi Hu, Ross M Kedl, Erika L Lasda, Jay R Hesselberth, MacLean Paul, Kimberley D Bruce, Gwendalyn J Randolph, Richard D Schulick, Yuwen Zhu

**Author notes:** Corresponding author. (Y.Z).

## Abstract

The lymphatic system plays a central role in lipid absorption, which transports chylomicrons from the small intestine to the circulation. However, the molecular mechanism by which chylomicrons get into the intestinal lymphatics is unknown. Here we demonstrated that GPR182, a receptor in lymphatic endothelial cells (LECs), mediates dietary fat absorption. GPR182 knockout mice are resistant to dietary-induced obesity. GPR182 ablation in mice leads to poor lipid absorption and thereby a delay in growth during development. GPR182 binds and endocytoses lipoproteins broadly. Mechanistically, loss of GPR182 prevents chylomicrons from entering the lacteal lumen of the small intestine. GPR182 blockage with a monoclonal antibody (mAb) protects mice from dietary induced obesity. Together, our study identifies GPR182 as a lipoprotein receptor that mediates dietary fat absorption.

## Introduction

Lipid transport, facilitated by lipoproteins and their corresponding receptors, plays a pivotal role in distributing essential nutrients and endogenously synthesized lipids to various tissues (*1*). Dysregulation of these processes, or lipid disorders, are major risk factors for metabolic diseases, which include, but are not limited to, atherosclerosis, fatty liver disease, type 2 diabetes, and neurodegenerative disorders (*2, 3*). Unlike nutrients such as amino acids and simple sugars that directly enter the bloodstream, lipid absorption involves the lymphatic system (*4, 5*). In the small intestine, dietary fatty acids are repackaged into particles called chylomicrons before entering the lymphatic structure known as lacteals (*6*). Despite its critical importance as a necessary step in fat absorption, the process through which chylomicrons enter into lacteals is poorly understood (*7*). The prevailing view assumes that chylomicrons reach the lacteal lumen through open cell-to-cell junctions between lymphatic endothelial cells (LECs), known as “button junctions,” which allow the passage of large cargoes like chylomicrons (*7, 8*). The conversion of intestinal lacteals from button-like to zipper-like junction leads to defects in fat absorption (*8, 9*). However, a receptor-mediated transcytosis mechanism for chylomicron transport has not yet been excluded (*10–12*).

GPR182 is a newly characterized atypical chemokine receptor (ACKR) of the G protein-coupled receptor (GPCR) superfamily (*13–15*). GPR182 is primarily expressed by microvascular, sinusoidal, and lymphatic endothelial cells, and GPR182 interacts with chemokines broadly via the glycosaminoglycan (GAG)-binding motif (*13, 14, 16*). In this study, we found that GPR182 interacts with lipoproteins broadly and ablation of GPR182 in mice leads to resistance to high fat diet (HFD)-induced obesity. Our study further supports that GPR182 serves as a receptor to mediate dietary fat absorption in the small intestine.

## Results

### GPR182^-/-^ mice are protected from dietary-induced obesity

Adult young GPR182^-/-^ mice on a C57BL/6J background grow healthy and breed normally on regular diets, consistent with previous reports (*13, 17*). However, we observed that GPR182^-/-^ mice weighed consistently less than their wild type (WT) littermates as they aged, with the difference between male mice being more prominent (**fig. S1A**). Male GPR182^-/-^ mice over 5-months-old became visibly slimmer in body shape (**fig. S1B)**. Examination of body composition via Magnetic Resonance Imaging (MRI) confirmed that male GPR182^-/-^ mice had a selective decrease in fat mass than control WT mice (**fig. S1C**). Hepatic steatosis observed in male WT mice at 15-month-old was absent in the livers of age-matched GPR182^-/-^ mice (**fig. S1D**).

The lymphatic system is known to mediate fat absorption and GPR182 is primarily expressed in LECs (*13, 14*). The observation of less weight in adult GPR182^-/-^ mice propelled us to carefully examine the role of GPR182 in dietary fat absorption. We monitored weight gain for young adult WT and GPR182^-/-^ mice over a 16-week period of standard chow diet or high-fat diet (HFD). At 7-week-old, GPR182^-/-^ male mice under chow diet weighed slightly less than age-matched WT controls (**Fig. 1A**). Over 16-week of chow diet, GPR182^-/-^ mice gained weight at a slower pace than WT controls, (**Fig. 1A-1C**). The most striking observation was the substantial resistance of GPR182^-/-^ mice to HFD-induced obesity (**Fig. 1A, 1B**). Upon 16 weeks of HFD feeding, male GPR182^-/-^ mice gained about 13 grams in body weight, while WT controls doubled their body weight (gained over 26 g/mouse, **Fig. 1C**). Consequently, the fat mass in GPR182^-/-^ mice was far less than that of WT controls (**Fig.1D**, **fig. S1E**). White adipose tissues (WATs) in GPR182^-/-^ mice were much smaller and weighed proportionally less than those from WT mice (**Fig. 1D**). There was no difference in brown adipose tissues (BATs) between GPR182^-/-^ and WT mice (**Fig. 1D**). Histological analysis confirmed adipocyte hypertrophy in WT mice upon 16-week of HFD feeding, and adipocytes in HFD-fed GPR182^-/-^ mice were markedly smaller (**Fig. 1E**). Livers from GPR182^-/-^ mice under HFD appeared normal and weighed less than half of those from WT counterparts (**Fig. 1F**). Triacylglycerol (TAG) in GPR182^-/-^ livers was only about 50% of those from WT mice (**Fig. 1G**). H&E and Oil Red O staining further confirmed that GPR182^-/-^ livers contained notably smaller and fewer lipid droplets than those of WT mice (**Fig. 1H**).

**Figure 1.**
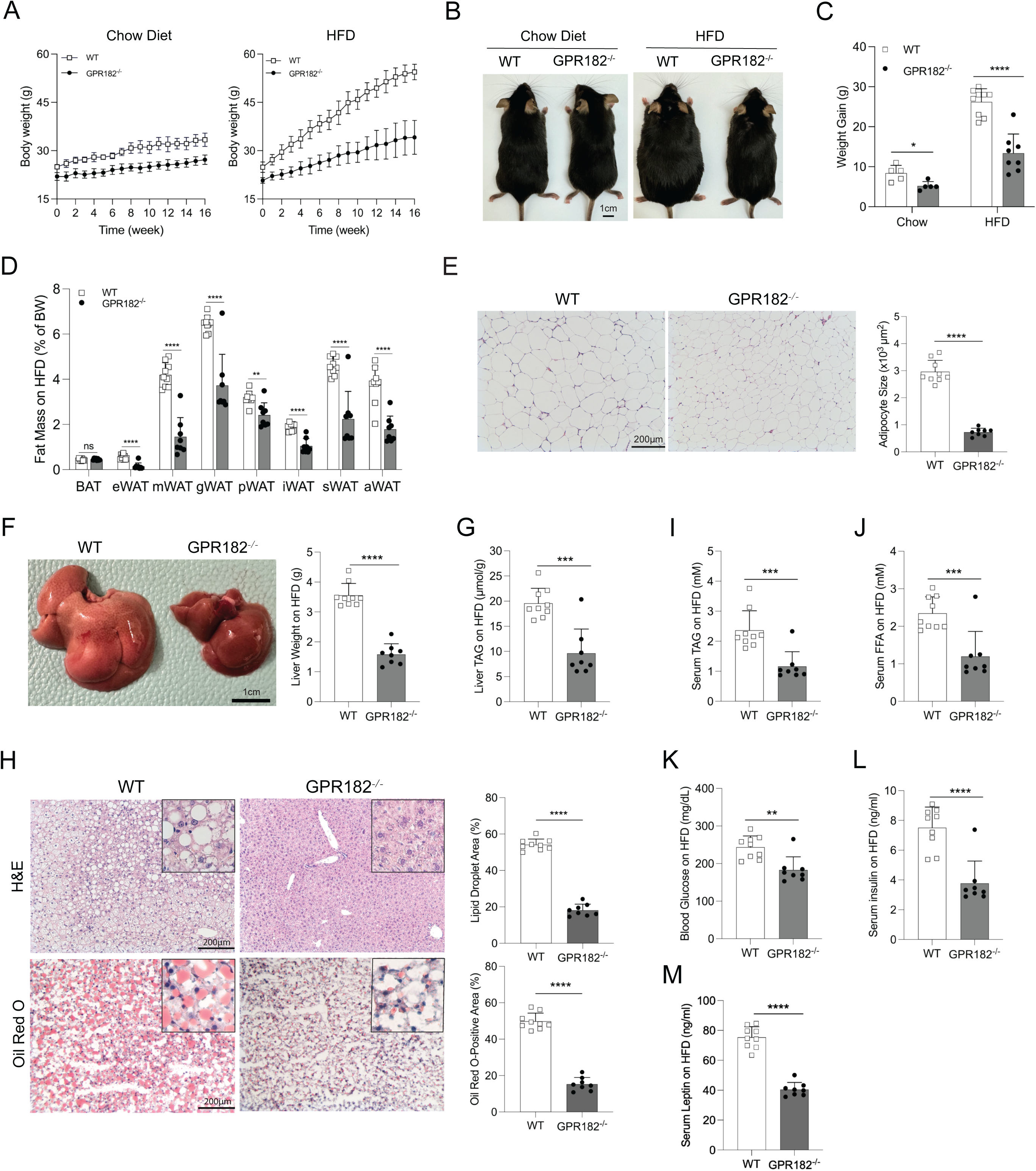
GPR182^-/-^ mice are resistant to diet induced obesity. 7-week-old male WT and GPR182^-/-^ mice were fed on chow diet or HFD for 16 weeks. (A) Body weight was followed weekly. (B) Representative images of mice upon 16 weeks of feeding were shown and (C) body weight gain was recorded. (D) Fat percentages of WT and GPR182^-/-^ mice fed on HFD were calculated. BAT, brown adipose tissue; eWAT, epicardial WAT; mWAT, mesenteric WAT; gWAT, gonadal WAT; pWAT, perirenal WAT; iWAT, inguinal WAT; sWAT, subcutaneous WAT; aWAT, axillary WAT. (E) H&E staining of gWAT from GPR182^-/-^ and control mice on HFD. Adipocyte sizes were determined in the right panel. (F) Representative images of livers from GPR182^-/-^ and WT mice on HFD were shown. Liver weight was calculated in the right panel. (G) H&E and Oil Red O staining of liver tissues from GPR182^-/-^ and WT control mice were recorded. Liver TAG (H), serum levels of TAG (I), FFA (J), glucose (K), insulin (L), and leptin (M) in GPR182^-/-^ and control WT mice on HFD were quantified. Each symbol represents one mouse. Chow diet: n=5; HFD: n=8-9.

Serum TAG and free fatty acid (FFA) from fasting GPR182^-/-^ mice after 16-week of HFD feeding were only about 50% of those in WT control (**Fig. 1I, 1J**). Further, GPR182^-/-^ mice exhibited significantly lower concentrations of circulating glucose, insulin, and leptin (**Fig. 1K-1M**). Consequently, GPR182^-/-^ mice on HFD were more resistant to glucose intolerance compared to WT counterparts (**fig. S1F**). Together, our results strongly support that GPR182 ablation protects mice from diet-induced obesity and its related disorders.

### GPR182 ablation prevents lipid absorption in small intestine

To better understand how GPR182^-/-^ mice were protected from HFD-induced obesity, we performed energy balance assessment for mice under two weeks of HFD. GPR182^-/-^ mice on HFD had the same daily food intake as their WT counterparts (**Fig. 2A**), whereas they gained less weight in a 2-week span of HFD feeding (**fig. S2A**). The energy expenditure under the light circadian phase was similar between GPR182^-/-^ and WT control mice, while GPR182^-/-^ mice tended to consume slightly more energy during the dark circadian phase (**fig. S2B**). This is consistent with more O_2_ consumption and CO_2_ production for GPR182^-/-^ mice during the dark phase **(fig. S2C)**. However, this minor change of energy consumption could not explain the far less weight gain for GPR182^-/-^ mice (**fig. S2A**). We found that GPR182^-/-^ mice produced significantly more feces, even though their food consumptions were the same (**Fig. 2B**). Feces produced by GPR182^-/-^ mice contained 23% more residual lipids than those from the WT control group (**Fig. 2C**), suggesting that loss of GPR182 reduces intestinal lipid uptake. As a result, GPR182^-/-^ mice on 2-week of HFD showed lipid droplet accumulation in the small intestine which was not found in control WT mice (**Fig. 2D**). This implies that GPR182^-/-^ mice have poor lipid absorption which leads to the increase of lipid excretion via feces.

**Figure 2.**
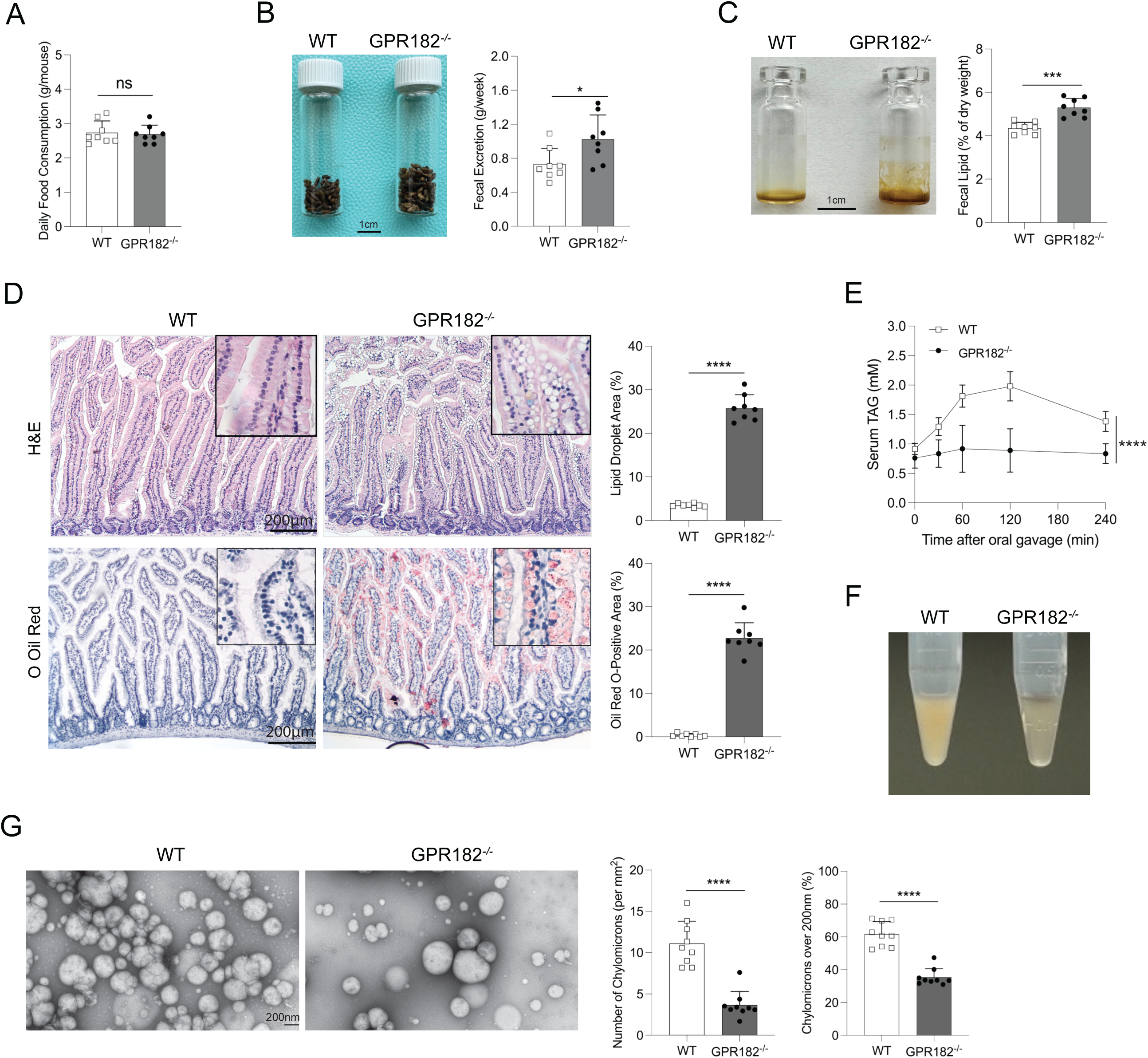
GPR182 ablation limits lipid absorption in small intestine. (A-C) Age-matched adult WT and GPR182^-/-^ male mice were fed with 2-week of HFD. Food consumption (A), feces production (B), and fecal lipids (C) were measured. n=8. (D) Jejunums from WT and GPR182^-/-^ mice on two weeks’ HFD were examined by H&E and Oil O Red staining. n=8. (E-G) Oral gavage of olive oil in WT and GPR182^-/-^ mice upon fasting. (E) Serum TAG in GPR182^-/-^ and WT mice were quantified over a 4-hour period upon olive oil gavage. n=8. (F) Representative images of serum from WT and GPR182^-/-^ mice 2 hours after oil gavage. (G) Transmission electron microscopy (TEM) analysis of mesenteric lymph fluid for chylomicrons from WT and GPR182^-/-^ mice 30 min after oil gavage. The number and size of chylomicrons were quantified. Each symbol represents one field. n=9. 3 mice in each group.

We evaluated dietary lipid absorption in fasting mice by olive oil oral gavage. Tyloxapol was injected simultaneously to inhibit lipoprotein catabolism. Upon overnight fasting, GPR182^-/-^ mice exhibited slightly lower serum TAG than WT mice (**Fig. 2E**). As expected, serum TAG in WT mice spiked high right after oil gavage. In contrast, GPR182^-/-^ mice maintained lower levels of serum TAG throughout the assessment (**Fig. 2E**). Consistently, serum from WT mice 2 hours after oil gavage was creamy white, whereas GPR182^-/-^ serum remained clear (**Fig. 2F**). We extracted lymph fluid from mesenteric lymphatics 30 min after oil gavage and performed transmission electron microscopy (TEM). Lymph liquid from GPR182^-/-^ mice contained considerably fewer chylomicron particles than that of WT mice, and chylomicron particles from GPR182^-/-^ mice were generally smaller (**Fig. 2G**). Consistently, we observed weaker fluorescence signal in GPR182^-/-^ mesenteric lymph ducts when we administered mice orally with a fluorescently labeled fatty acid (BODIPY FL C_16_) (**fig. S2D**). Histological analysis did not reveal any structural change in intestinal villus from GPR182^-/-^ mice (**fig. S2E**). In addition, the draining function of intestinal lymphatics between WT and GPR182^-/-^ mice was comparable when we assessed by injection of FITC-Dextran into the Peyer’s patch (**fig. S2F**). Therefore, GPR182 ablation in mice leads to a selective defect in dietary lipid absorption.

### GPR182 interacts with lipoproteins broadly

Our previous study revealed that GPR182 is an ACKR that interacts with chemokines via the GAG-binding motif (*13*). Many apolipoproteins or lipoproteins are known to interact with GAGs (*18, 19*). Given what we observed of GPR182^-/-^ mice in diet-induced obesity, we hypothesized that GPR182 is a receptor on lymphatics to mediate lipoprotein transport. First, we confirmed that apolipoprotein B-100 (ApoB-100) (**Fig. 3A**) and apolipoprotein E (ApoE) (**Fig. 3B**), two major ligand-binding apolipoproteins with known GAG-binding capacities (*20*), bound to human GPR182^+^ 293T cells by flow cytometry. Inclusion a monoclonal antibody (mAb) against GPR182 (clone 1A5) (**fig. S3A**) markedly reduced the bindings (**Fig. 3A, 3B**), demonstrating the specific interaction between GPR182 and these two apolipoproteins.

**Figure 3.**
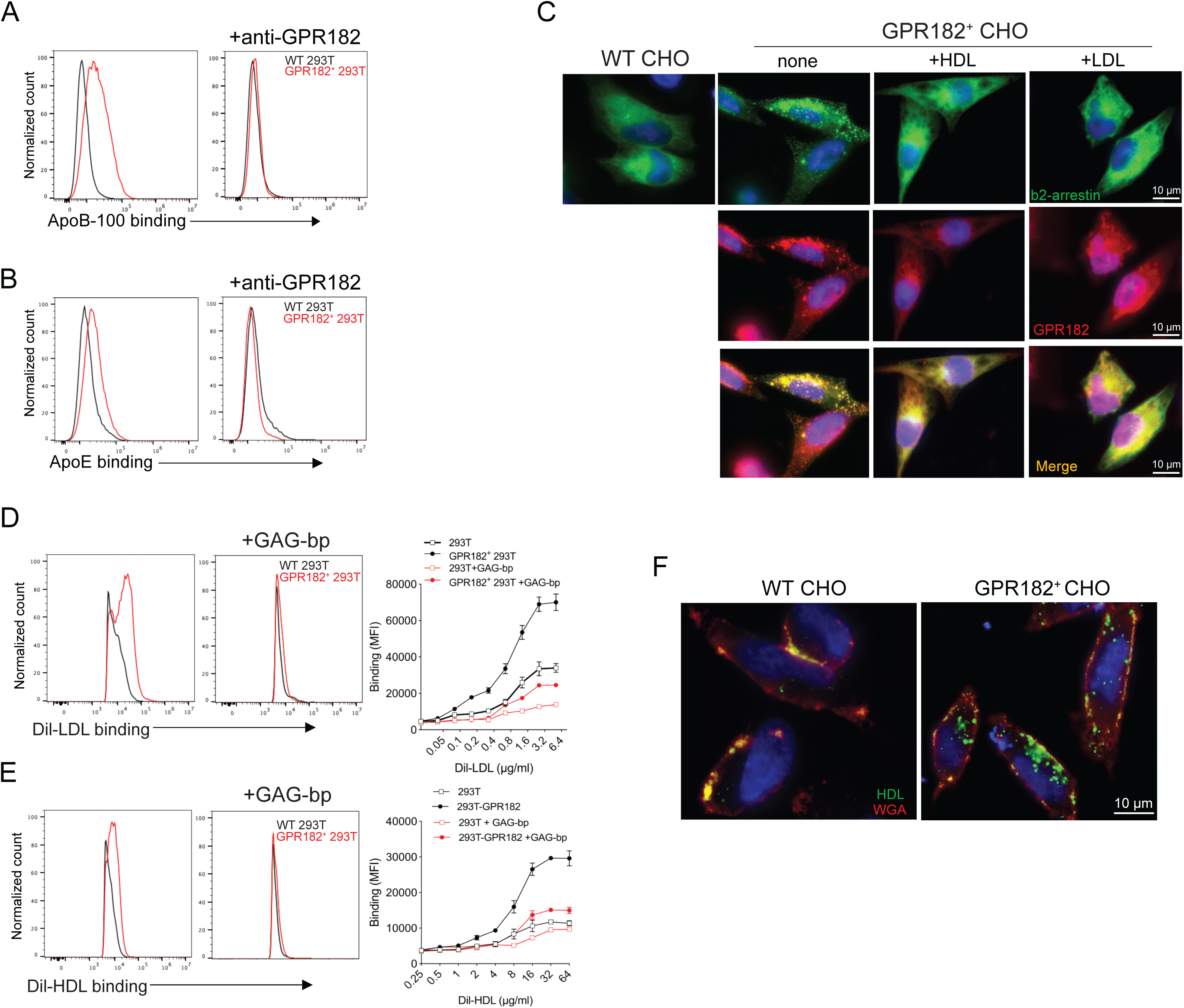
GPR182 interacts with lipoproteins broadly. (A, B) GPR182^+^ HEK293T and control cells were examined for protein ApoB-100 (A) and ApoE (B) binding by flow cytometry. GPR182 mAb was added to assess its blocking capacity. (C) GPR182^+^ CHO and control CHO cells were transfected to express β2-arrestin-eGFP. GPR182^+^ CHO cells were incubated with HDL or LDL at 37 degrees for one min before immunofluorescence staining to locate β2-arrestin and GPR182. (D) GPR182^+^ HEK293T and control cells were examined for Dil-LDL binding by flow cytometry. GAG-bp was added to assess its blocking capacity. (E) GPR182^+^ HEK293T and control cells were examined for Dil-HDL binding by flow cytometry. GAG-bp was added to assess its blocking capacity. (F) GPR182^+^ CHO and control CHO cells were incubated with Dio-HDL at 37 degrees for 5 min. Cells were fixed and co-stained with wheat-germ agglutinin (WGA) before imaged by confocal microscopy.

GPR182 is a newly-characterized ACKR that constitutively associates with β2-arrestin (*13–15*). In a NanoBit assay, co-transfection of GPR182 and β2-arrestin in HEK293T cells triggered a strong interaction signal (*13*). The addition of human serum was able to effectively abolish the signal (**fig. S3B**). Similarly, all purified serum lipoproteins inhibited the GPR182/β2-arrestin interaction in a dose-dependent manner (**fig. S3C**). Among them, very low-density lipoprotein (VLDL) and chylomicron were the two more effective lipoproteins to disrupt the GPR182/β2-arrestin association, in comparison with high-density lipoprotein (HDL) and low-density lipoprotein (LDL). We expressed β2-arrestin-eGFP in GPR182^+^ and control CHO cells to better visualize the interaction between β2-arrestin and GPR182. β2-arrestin was evenly distributed within the cytoplasm of WT CHO cells, whereas in GPR182^+^ CHO cells β2-arrestin tended to associate with GPR182 to aggregate. Interestingly, the addition of HDL or LDL in GPR182^+^ CHO cells was able to effectively disrupt the aggregation between GPR182 and β2-arrestin (**Fig. 3C**).

We directly assessed whether lipoproteins from human serum interact with GPR182. At 4 °C, Dil-LDL bound to GPR182^+^ HEK293T cells far stronger than WT HEK293T cells; the addition of a generic GAG-binding peptide (GAG-bp) (*13*) was able to eliminate the binding (**Fig. 3D**). Similarly, Dil-HDL bound to GPR182^+^ CHO cells, which could be competed off by the same GAG-bp (**Fig. 3E**). All four serum lipoproteins effectively competed with chemokine CXCL10 for GPR182 binding, suggesting that chemokines and lipoproteins share a similar binding site on GPR182 (**fig. S3D**). The GPR182 mAb clone 1A5, which blocks the CXCL10/GPR182 interaction (**fig. S3E**), was also able to inhibit Dil-HDL binding effectively (**fig. S3F**). When we incubated Dio-HDL with CHO cells at 37 °C, GPR182^+^ CHO, not WT CHO cells, were able to endocytose HDL efficiently (**Fig. 3F**). Taken together, our results supported GPR182 as a common receptor for lipoproteins.

### GPR182 regulates lipoprotein homeostasis

Besides LECs, GPR182 is expressed by microvascular and sinusoidal ECs in multiple organs (*14, 21, 22*). The broad expression of GPR182 led us to assess the possible impact of GPR182 ablation on lipoprotein homeostasis. GPR182^-/-^ mice on regular chow diet had significantly higher total serum cholesterol than age-matched WT mice (**Table S1**). Cholesteryl ester, but not free cholesterol, was increased in the serum of GPR182^-/-^ mice. In contrast, serum levels of TAG and FFA were slightly but significantly reduced in GPR182^-/-^ mice (**Table S1**). All these changes were consistently found in both female and male GPR182^-/-^ mice at different ages (**Table S1**). Fast protein liquid chromatography (FPLC) analysis further revealed a selective increase of serum HDL-cholesterol in GPR182^-/-^ mice (**fig. S4A**). Mass spectrometry analysis of serum proteins from GPR182^-/-^ mice revealed that multiple serum apolipoproteins were increased, including Apolipoprotein C-IV, Apolipoprotein D, Apolipoprotein A-II, Apolipoprotein B-100, and Apolipoprotein C-I (**fig. S4B**) (*23*). Our results thus implicate that GPR182 contributes to cholesterol transport *in vivo*.

### GPR182 on lacteals mediates chylomicron transportation

We further confirmed the involvement of GPR182 in lipid absorption by analyzing newborns during the lactation period. GPR182^-/-^ pups were visually indistinguishable from WT littermates (**fig. S5A**). However, they weighed slightly but significantly less than their WT littermates throughout the course of lactation (**Fig. 4A**). Collecting mesenteric lymphatics from 7-day-old (P6) GPR182^-/-^ newborns were pale or transparent whereas those from WT counterparts appeared milky white (**Fig. 4B**). Quantification of lymphatic collectors in P6 pups revealed that GPR182^-/-^ pups contained far fewer collecting lacteals filled with white chylomicrons than WT controls (**Fig. 4C**). TEM analysis confirmed a remarkable reduction of chylomicrons in mesenteric lymph fluid from GPR182^-/-^ newborns (**Fig. 4D**).

**Figure 4.**
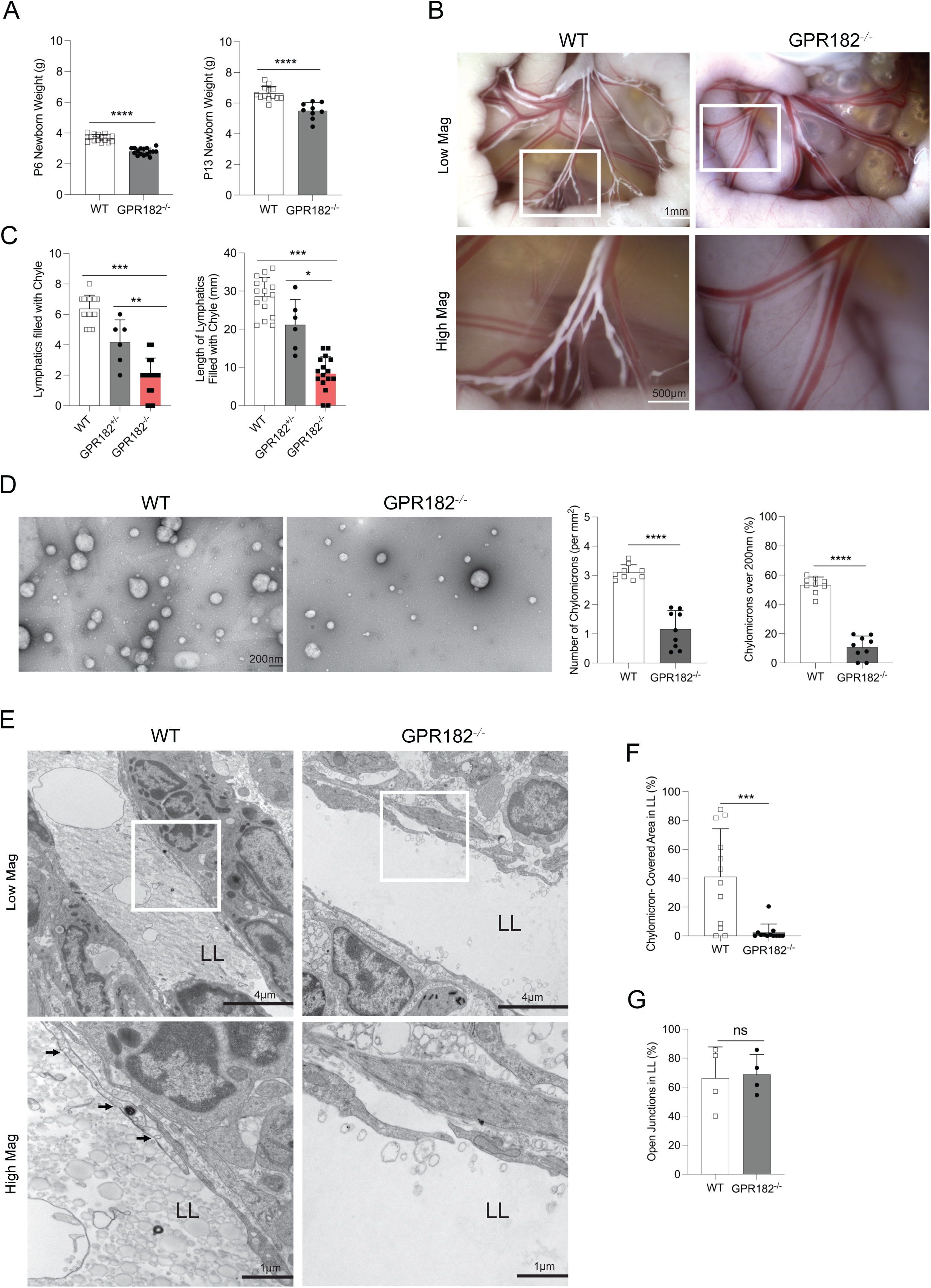
Limited chylomicron transport in GPR182^-/-^ lacteals. (A) Body weights for GPR182^-/-^ and littermates at P6 (n=16) and P13 (n=9, 13) were measured. (B, C) P6 newborns of GPR182^-/-^ and its littermates were examined for mesenteric lymphatics. Representative images (B) and quantifications (C) of lymphatics filled with chylomicrons were recorded. Each symbol represents one mouse. (D) TEM analysis of mesenteric lymph fluid for chylomicrons from P6 newborns of WT and GPR182^-/-^ mice. The number and size of chylomicrons were quantified. Each symbol represents one field (n=3 per group). (E) TEM analysis of intestinal villus from adult WT and GPR182^-/-^ mice after olive oil gavage. LL: lacteal lumen. Quantification of chylomicrons in intestinal lacteal lumens (F) and open lacteal junctions (G) via TEM in adult mice. Each symbol represents one lacteal lumen (F) or mouse (G), n=4 per group.

In the small intestine, GPR182 is exclusively expressed by Lyve1^+^ LECs (**fig. S5B**) (*14, 22*). We performed TEM to visualize chylomicron transport in lacteals. In adult mice upon oil gavage, lipid droplets or chylomicrons were abundantly present in enterocytes and the interstitial space between enterocytes and LECs. Few chylomicrons were present in the lacteal lumens of GPR182^-/-^ mice, which was in sharp contrast to WT lumens full of chylomicron particles (**Fig. 4E**). Quantification analysis further confirmed a sharp reduction of chylomicrons within the lacteal lumens of GPR182^-/-^ mice (**Fig. 4F**). In newborn pups under breastfeeding, lacteal lumens from WT newborns were abundantly filled with chylomicrons while chylomicrons were scarce in lacteals from GPR182^-/-^ pups (**fig. S5C, S5D**). There was no structural alteration in junction openness of lacteal lumens in GPR182^-/-^ mice (**Fig. 4G**, **fig. S5E**). VE-cadherin staining in intestinal lacteals also revealed a similar button-like/zipper-like pattern between adult GPR182^-/-^ mice and control WT mice (**fig. S5F**). Together, our results support that GPR182 on lacteals is required for chylomicron transport in the small intestine.

### GPR182 blockade limits dietary lipid absorption to treat HFD-induced obesity

We assessed whether blockage of GPR182 can limit dietary lipid absorption and therefore reduce HFD-induced obesity. We used homogeneous human GPR182 knock-in (hGPR182-KI) mice for this study because our GPR182 mAb (clone 11C7) recognizes human, but not mouse, GPR182. This mAb was able to effectively inhibit HDL binding (**fig. S6A**) and lower serum TAG in hGPR182-KI mice upon olive oil oral gavage (**fig. S6B**). We administrated hGPR182-KI mice with 11C7 or control mIgG1 weekly during a 16-week span of HFD challenge. Anti-GPR182 mAb did not have an instant impact on weight gain in response to HFD. Only after 4-week of HFD feeding, we began to observe weight gain difference between these two mouse groups (**Fig. 5A**). After 16-week of HFD, the average body weight of control mice was 48 grams while mice treated with GPR182 mAb was about 41 grams (**Fig. 5B, 5C**). The reduced weight gain in GPR182 mAb-treated mice was solely due to less fat mass as the two groups had the same weight of the lean mass (**Fig. 5D**). Consistently, fat tissues from mice treated with 11C7 weighed less and the adipocytes were smaller (**Fig. 5E, 5F, fig. S6C, S6D**). The treatment of 11C7 in HFD-challenged mice significantly reduced liver weight and alleviated the symptoms of fatty liver (**Fig. 5G, 5H**). Furthermore, mice treated with anti-GPR182 produced more feces, which contained higher levels of lipid residues (**fig. S6E, S6F**). All these findings were consistent with what we observed in GPR182^-/-^ mice under HFD challenging.

**Figure 5.**
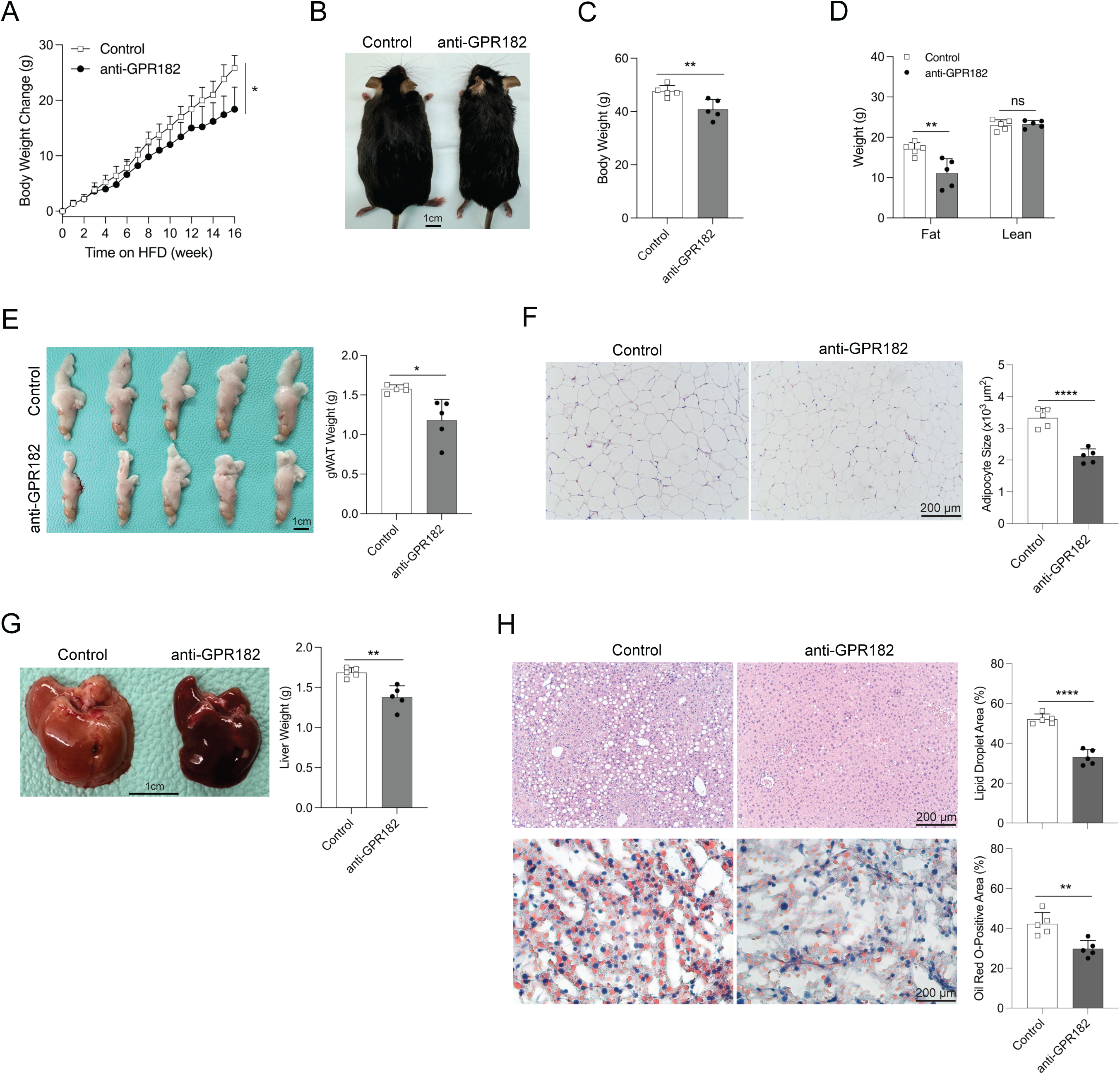
GPR182 blockade alleviates HFD-induced obesity. (A-H) 7-week-old male hGPR182-KI mice were fed on HFD for 16 weeks. Mice were treated with control or anti-hGPR182 mAb weekly at the beginning of HFD feeding. n=11,12. (A) Body weight gain was followed weekly. (B) Representative images of hGPR182-KI mice upon 16 weeks of HFD feeding were shown and (C) body weight was recorded. (D) Fat and lean masses of mice were determined by MRI. (E) Representative images of gWATs were shown. gWAT weight was determined in the right panel. (F) H&E staining of gWATs was shown. Adipocyte sizes were determined in the right panel. (G) Representative images of livers from antibody-treated mice on HFD were shown. Liver weight was calculated in the right panel. (H) H&E and Oil Red O staining of liver tissues from antibody-treated mice were recorded and quantified. n=5.

We further tested whether GPR182 blockade can be a therapeutic approach for diet induced obesity. Diet-induced obese hGPR182-KI mice were treated twice a week with control or GPR182 mAb and continued with HFD feeding. After eight weeks of treatment, anti-GPR182 selectively reduced total fat mass while it had no impact on lean mass (**fig. S7A, S7B**). Consistently, mice treated with GPR182 mAb had significantly smaller gonadal WAT (**fig. S7C**) and mesenteric WAT (**fig. S7D**). GPR182 mAb-treated mice had greatly improved fatty livers (**fig. S7E, S7F**). Together, our study supports GPR182 blockade as a viable approach to reduce obesity and its related malfunctions.

## Discussion

We demonstrated that GPR182 serves as a lipoprotein receptor that mediates fat absorption in the small intestine. This finding is based on our studies of GPR182^-/-^ mice and the usage of a GPR182 blocking mAb. Our findings thus provide direct molecular evidence to support a receptor-mediated transcytosis mechanism for dietary fat absorption.

Lacteals in the small intestine are known to mediate lipid absorption in the form of chylomicrons. Chylomicrons have been thought to be transported paracellularly as the discontinuous, button-like junctions in lymphatic capillaries allow chylomicrons to cross (*24*). However, few large open junctions in lacteals observed by early TEM imaging would argue against its role as the primary route for chylomicron transport into the lacteals (*10, 25*). Further, it was reported that, in a vitro model, the transport of lipid across the lymphatic endothelium occurs via an ATP-dependent, transcellular route (*11*). Our results support receptor-mediated transcytosis as the primary mechanism for chylomicron uptake in the small intestine. Loss of GPR182 leads to a dramatic reduction of chylomicrons crossing into lacteal lumens in the small intestine. As a result, GPR182^-/-^ mice under HFD display lipid droplet accumulation in the small intestine. GPR182^-/-^ pups exhibit retarded weight gain during breastfeeding and adult GPR182^-/-^ mice are resistant to HFD-induced obesity. This protection is not caused by a defect in lymphatic function as lymphatic draining in GPR182^-/-^ mice appears to be normal (*13*). Further, the application of a GPR182 blocking mAb in the HFD-induced obesity model recapitulates what we observed in GPR182^-/-^ mice (**Fig. 5**). All these findings support a major role of GPR182 in mediating dietary lipid absorption. On the other hand, our study does not negate the involvement of free chylomicron transport through open cell-to-cell junctions between LECs within the lacteals. We still observed some chylomicron particles in lacteal lumens of GPR182^-/-^ mice upon olive oil gavage. In addition, some mesenteric connecting lymphatics from GPR182^-/-^ newborns were still filled with chylomicrons. The respective contribution in relation to these two different processes for fat absorption would require careful comparison in different biological conditions.

GPR182 belongs to a group of GPCRs known as ACKRs which primarily function as chemokine scavengers (*26*). However, known ligands for ACKRs are not exclusively chemokines. For example, ACKR3 is a receptor for adrenomedullin (*27*) and a broad scavenger of opioid peptides (*28, 29*). CCRL2 and GPR1, two new members of ACKRs, bind chemerin, a non-chemokine attractant (*26*). Our previous study identifies GPR182 as a pattern-recognition receptor for GAG-binding peptides (*13*). Similar to chemokines, apolipoproteins such as ApoB and ApoE are known to be GAG-interacting proteins (*18, 30*). Here we were able to confirm their GPR182 binding and found that lipoproteins and chemokines share a similar binding site on GPR182. Compared with most chemokines, lipoproteins have relatively weaker binding to GPR182 (*13, 14, 16*). This is reasonable as lipoproteins are far abundant than chemokines in the body for GPR182 to interact with. Consistently, a recent report revealed that GPR182 interacts with many non-chemokine proteins (*31*). In a high-throughput screening assay, GPR182 has been screened out as one of three candidates that are potential binding receptors for LDL (*32*). Therefore, our characterization of GPR182 as a lipoprotein receptor is not completely surprising.

The broad expression of GPR182 in the lymphatic system would support its involvement in both endogenous lipid transport and exogenous fat absorption. Although GPR182 serves as a receptor for all lipoproteins, GPR182^-/-^ mice on regular chow diet are generally normal and only exhibit higher circulating HDL in peripheral blood. Supporting that, in a genome-wide association study (GWAS), a GPR182 gene variant (rs58298943-C) is significantly associated with high serum HDL (*p*-value at 4x10^-9^) (*33*). The presence of other specialized lipoprotein receptors could play a complementary role to compensate for the loss of GPR182 (*34, 35*). It would be interesting to further understand how GPR182 coordinates with other known lipoprotein receptors to regulate lipoprotein transport and metabolism in various pathological conditions.

In conclusion, our studies identify a lipoprotein receptor in GPR182 that mediates lipid absorption in the small intestine. Agents to prevent GPR182 from lipoprotein transport can be an interesting approach of treating obesity and other disorders related to lipid metabolism.

## Materials and methods

### Mice and antibody

GPR182^-/-^ mice were generated by crossing GPR182^lacZ/lacZ^ mice (*13*) with CMV-Cre mice from the Jackson Laboratory (Strain #:006054). Genotyping for GPR182^-/-^ mice was performed with the following primers: Forward: GCTACCATTACCAGTTGGTCTGGTGTC; Reverse: AGAGAAAGGTCATCTGTGAGGAGGC. Human GPR182-knockin (hGPR182-KI) mice were generated by Leveragen Inc. with the CRISPR/Cas9 technology, by replacing the coding sequence of mouse GPR182 with human GPR182 through homologous recombination. All mice were backcrossed to C57BL/6J background and were housed in a pathogen-free vivarium with a 14-hour light/10-hour dark cycle at the University of Colorado Anschutz Medical Campus (CU AMC). C57BL/6J mice purchased from the Jackson laboratory were housed at the CU AMC animal facility for at least two weeks before usage. Except the HFD model, both male and female mice were used for experiments.

Monoclonal antibodies (mAbs) against human GPR182 were generated by hybridoma fusion of splenocytes from immunized GPR182^-/-^ mice. Antibody in culture medium was purified with a Protein G HP Column (Cytiva HiTrap™). The specificity of hGPR182 mAbs were confirmed by their specific binding to GPR182-transfected cells by flow cytometry. The blocking capacity was further assessed by their effect in blocking the binding between GPR182^+^ cells and CXCL10-AF647 (ALMAC) and Dil-HDL (Kalen Biomedical, LLC).

### HFD model

Male wild type (WT) and GPR182^-/-^ mice at 7-10 weeks old were fed with rodent diet with 58 kcal% fat and sucrose (Research Diets: D12331) for 16 weeks. 42 g/L of carbohydrates was mixed in drinking water at a ratio of 55% fructose and 45% sucrose by weight (Sigma-Aldrich, St. Louis, MO). Mice had ad libitum access to food and water and were weighed weekly. For GPR182 mAb administration, male hGPR182-KI mice at 8-week-old were injected intraperitoneally with control mIgG1 (BioXcell) or anti-GPR182 mAb at 300µg/mouse. Mouse weight was measured weekly. All animal experiments were approved by the Institutional Animal Care and Use Committee at CU AMC.

### Energy Balance Assessment

Energy balance assessment for GPR182^-/-^ and control WT mice under HFD was conducted at the Energy Balance Assessment Core of Colorado Nutrition Obesity Research Center. Total energy expenditure (TEE), resting energy expenditure (REE) and non-resting energy expenditure (NREE) were measured in Oxymax chambers in comprehensive lab animal monitoring system (CLAMS) (Columbus Instruments, Columbus, OH). Mice were placed individually in metabolic chambers with an animal activity meter (Opto-Max, Columbus Instruments, Columbus, OH) for two weeks. Mice had ad libitum access to food and water. Metabolic chambers were maintained at 23 °C with 12 hours light/dark cycles. Metabolic rate (MR) was calculated from gas exchange measurements acquired every 18 min using the Weir equation (*36*). Then MR was averaged and extrapolated over 24 hours to estimate TEE. Before and after the energy balance assessment, whole-body fat and lean composition were measured by Echo MRI (Echo Medical Systems, Houston, TX).

### Lipoprotein, TAG, lipids, and FFA quantification

Blood samples were centrifuged at 1000 × g for 10 min at 4 °C to extract the plasma. The concentrations of cholesterol, TAG, FFA, and glucose in plasma samples were measured following the manufacturer’s instructions of colorimetric assay kits (Merck-Millipore). The levels of leptin and insulin in plasma samples were quantified with ELISA kits (Merck-Millipore).

Lipids in mouse tissues were extracted using the Folch method (*37*). Fecal lipids were extracted using a modified Folch method (*38*). Briefly, samples were homogenized in chloroform in methanol (2:1 by volume), and centrifuged at 1000 × g for 10 min. The supernatants were collected, dried and weighted. Finally, the extracts were re-dissolved in ethanol to measure lipids with the assay kits as described above.

Plasma samples (200 µL) were chromatographed via fast protein liquid chromatography (FPLC) using two Superose 6 columns in series as previously reported (*39*). During size exclusion, the absorbance of the eluted samples was measured at 280 nm, allowing identification of each lipoprotein class via protein content. Fractions containing either VLDL, LDL, or HDL were pooled, and cholesterol was measured using a commercially available kit (Cayman Chemical Company, Ann Arbor, MI, USA) following manufacturer’s instructions.

### Oil gavage assay

After 12 hours of fasting and 30 min of tyloxapol pretreatment intraperitoneally (500 mg/kg of body weight, Merck-Millipore), GPR182^-/-^ and control WT mice at the same age were given with olive oil by oral gavage (10 μL/g of body weight). Blood samples were collected via lateral tail veins before (time 0) and at 30, 60, 120 and 240 min after oil gavage. Levels of TAG in plasma samples were measured as described above. In some experiments, mice were pre-treated with GPR182 mAb at 300µg/mouse before oil gavage. Liver and intestine samples were collected 2 hours post oil gavage for H&E and Oil Red O staining. BODIPY FL C_16_ (Thermo Fisher, D3821) was added to olive oil to make final concentration at 0.4 μg/μL. After 6 hours of fasting, 2-week old GPR182^-/-^ and control WT mice were administered orally with diluted BODIPY FL C_16_ at 50 μL per mouse. Mice were euthanized 2 hours after the gavage. The whole intestine along with mesenteric lymphatics were put in 35mm glass bottom culture dish and imaged with Nikon Eclipse TE2000-E upright microscope.

### The NanoBit assay

HEK293T cells were co-transfected with two plasmids that encode for GPR182 tagged with LgBit and β2-arrestin tagged with SmBit (*13*). 24 hours later, cells were replated and incubated with lipoproteins at different concentrations. NanoGLO substrate was added right before luminescence measurement.

### Lipoprotein binding and endocytosis in vitro

For lipoprotein binding studies, both WT control and GPR182^+^ 293T cells were incubated with or without pretreatment of 1 mg/ml GAG-binding peptide at 4 °C for 15 min. Then cells were incubated with Dil-LDL or Dil-HDL (Kalen Biomedical, LLC) at 4 °C for 30 min. After cells were washed with cold PBS, flow cytometry was performed to assess the binding of lipoproteins to GPR182-expressing cells. FlowJo 11 was used to calculate mean fluorescence intensity (MFI) and Prism 9.0 was used to analyze non-linear regression. For lipoprotein endocytosis, WT CHO and GPR182^+^ CHO cells were serum-starved for 6 h before incubated with Dio-HDL at 37 °C for 5 min. Cells were stained with wheat germ agglutinin (WGA, Thermo Fischer) and spectral DAPI (4′,6-diamidino-2-phenylindole; Akoya Biosciences, Inc) before analysis. Images were taken by Nikon Eclipse TE2000-E upright microscope.

### Immunohistochemistry and immunofluorescent staining

For H&E staining, tissues were formalin-fixed and embedded in paraffin. The blocks were sectioned at 5 μm and stained with haematoxylin & eosin (H&E). For Oil Red O staining, tissues were frozen on dry ice in OCT embedding media. Then the blocks were sectioned at 8 μm and stained with Oil Red O, followed by counterstaining with hematoxylin. For Immunofluorescence staining, frozen blocks were sectioned at 8 μm and mounted on glass slides. The slides were then fixed in acetone and blocked with 2.5% goat serum before incubated with appropriate primary antibodies overnight at 4 °C. After washed three times with PBS, slides were stained with fluorophore dye conjugated secondary antibodies for 2 h and counterstained with DAPI for 10 min at room temperature. Slides were cleared and mounted with Fluoromount-G™ Mounting Medium (Thermo Fisher) before imaged with Nikon Eclipse TE2000-E upright microscope. Images were analyzed with SlideBook software (Version 6, Intelligent Imaging Inc). Sometimes images were taken by Olympus FV1000 FCS confocal laser scanning microscope and analyzed using ImageJ (National Institutes of Health).

### Transmission Electron Microscopy (TEM)

Jejunum tissues were dissected from mouse after transcardiac perfusion with saline followed by 4% paraformaldehyde and 1% glutaraldehyde in 0.1M sodium cacodylate buffer. Samples were then fixed overnight with 2.5% (v/v) glutaraldehyde in 0.1M sodium cacodylate buffer. After five rinses with 0.1M sodium cacodylate buffer, tissues were postfixed in 1% osmium tetroxide and 0.8% potassium ferrocyanide in 0.1M sodium cacodylate buffer for 1.5 hours. Tissue samples were rinsed five times in water and en bloc stained with 2% uranyl acetate in 50% ethanol for 2 hours. They were dehydrated with increasing concentrations of ethanol, transitioned into resin with propylene oxide, infiltrated with EMbed-812 resin and polymerized in a 60 °C oven overnight. Blocks were sectioned with a diamond knife (Diatome) on a UC7 ultramicrotome (Leica) and collected onto copper grids, post stained with 2% aqueous uranyl acetate and lead citrate. Images were acquired on a Tecnai T12 transmission electron microscope (Thermo Fisher) equipped with a LaB6 source at 120kV using a XR80 (8Mpix) camera (AMT).

### Negative Stain Electron Microscopy

Liquid was collected from mesenteric lymph nodes (mLNs) in mice and diluted at 1:50 in PBS. A 10 μL drop of sample was applied to a freshly glow-discharged 300 mesh formvar and carbon coated grid (Electron Microscopy Sciences) for 10 min and blotted with filter paper. The grid was then washed one time by applying 10 μL water on the grid and blotted with a Whatman filter paper. Finally, the grids were stained with 10 ml of 2% uranyl acetate solution for 3 min. After blotting, the grids were allowed to dry. Samples were imaged on a Thermo Fisher Tecnai G2 Biotwin TEM (ThermoFisher) at 120 kV with an AMT low mount NS15B sCMOS camera (AMT Imaging).

### Functional lymphatic flow

The intestinal lymphatic draining function was assessed by FITC-Dextran injection of a Peyer’s patch, as described previously (*40*). Briefly, the abdominal cavity of anesthetized mice was opened with a midline incision of the peritoneum to expose the intestine and mesentery. 20 μg of 2000 kDa FITC-Dextran (Sigma-Aldrich, FD2000S) was injected into a Peyer’s patch in the ileum. Blood samples were collected via lateral tail vein at 0, 5, 15, 30 and 60 min after injection. Mice were kept warm and the intestine and mesentery were kept moist throughout the process. Plasma fluorescence was measured using BioTek Synergy H1 Plate Reader. The excitation wavelength at 495 nm and the emission at 519 nm were used to detect the fluorescence.

### Data and statistical analysis

All the experiments were repeated independently. The data were presented as mean ± standard deviation (SD). Unpaired two-tailed Student’s t test was used to compare the means of two groups. One-way analysis of variance (ANOVA) was used to compare the means of more than two groups. Two-way ANOVA with Bonferroni’s correction for multiple comparison was used to compare groups over time by repeated measures. All *P* values less than 0.05 were deemed to be significant. ns, not significant. \**P*<0.05, \*\**P*<0.01, \*\*\**P*<0.001, \*\*\*\**P*<0.0001. GraphPad Prism 9.0 (GraphPad Software, Inc., La Jolla, CA, USA) was used for all statistical analysis and to generate figures.

## Acknowledgments

We thank Dr. Matthias Löhr at Karolinska Institutet for constructive advice about the role of lymphatics in fat absorption. We thank the Mass Spectrometry Proteomics Shared Resource at the University of Colorado Cancer Center, the Human Immune Monitoring Shared Resource, the Advanced Light Microscopy Core Facility, the Energy Balance Assessment Core at Colorado Nutrition Obesity Research Center, and the Electron Microscopy Core Facility at the University of Colorado Anschutz Medical Campus for resources and services used in this study.

## Author contributions

Z.S. and Y.Z. designed the study; Z.S. conducted experiments and acquired the data for most of the study; Y.Z., R.J.T., E.N.M., A.D., I.V., T.T., E.E., Y.S., Y.G., E.J.Y., E.L.L, J.H., J.R.H., K.D.B, and G.J.R. assisted with experiments and data acquirement; R.M.K. and P.M. provided with resource and assistance; Y.Z. and R.D.S. provided with resource and supervised the project; Y.Z. conceived and wrote the manuscript; all authors read and approved the final manuscript.

## Data and materials availability

All data associated with this study are present in the paper or the Supplementary Materials. GPR182^-/-^ mice, hGPR182-KI mice, and mAbs for GPR182 are available from University of Colorado Anschutz Medical Campus upon completion of a material transfer agreement.

## References and Notes

1. J. Boren, M. R. Taskinen, E. Bjornson, C. J. Packard, Metabolism of triglyceride-rich lipoproteins in health and dyslipidaemia. Nat Rev Cardiol 19, 577–592 (2022).

2. J. L. Goldstein, M. S. Brown, A century of cholesterol and coronaries: from plaques to genes to statins. Cell 161, 161–172 (2015).

3. P. Libby et al., Atherosclerosis. Nat Rev Dis Primers 5, 56 (2019).

4. J. Bernier-Latmani, T. V. Petrova, Intestinal lymphatic vasculature: structure, mechanisms and functions. Nat Rev Gastroenterol Hepatol 14, 510–526 (2017).

5. N. Escobedo, G. Oliver, The Lymphatic Vasculature: Its Role in Adipose Metabolism and Obesity. Cell Metab 26, 598–609 (2017).

6. G. J. Randolph, N. E. Miller, Lymphatic transport of high-density lipoproteins and chylomicrons. J Clin Invest 124, 929–935 (2014).

7. J. B. Dixon, Mechanisms of chylomicron uptake into lacteals. Ann N Y Acad Sci 1207 Suppl 1, E52–57 (2010).

8. F. Zhang et al., Lacteal junction zippering protects against diet-induced obesity. Science 361, 599–603 (2018).

9. S. H. Suh et al., Gut microbiota regulates lacteal integrity by inducing VEGF-C in intestinal villus macrophages. Embo Rep 20, (2019).

10. W. O. Dobbins, 3rd, Intestinal mucosal lacteal in transport of macromolecules and chylomicrons. Am J Clin Nutr 24, 77–90 (1971).

11. A. L. Reed, S. A. Rowson, J. B. Dixon, Demonstration of ATP-dependent, transcellular transport of lipid across the lymphatic endothelium using an in vitro model of the lacteal. Pharm Res 30, 3271–3280 (2013).

12. J. B. Dixon, S. Raghunathan, M. A. Swartz, A tissue-engineered model of the intestinal lacteal for evaluating lipid transport by lymphatics. Biotechnol Bioeng 103, 1224–1235 (2009).

13. R. J. Torphy et al., GPR182 limits antitumor immunity via chemokine scavenging in mouse melanoma models. Nat Commun 13, (2022).

14. A. Le Mercier et al., GPR182 is an endothelium-specific atypical chemokine receptor that maintains hematopoietic stem cell homeostasis. Proc Natl Acad Sci U S A 118, (2021).

15. S. Melgrati et al., GPR182 is a broadly scavenging atypical chemokine receptor influencing T-independent immunity. Front Immunol 14, 1242531 (2023).

16. R. Bonnavion, S. Liu, H. Kawase, K. A. Roquid, S. Offermanns, Large chemokine binding spectrum of human and mouse atypical chemokine receptor GPR182 (ACKR5). Front Pharmacol 14, 1297596 (2023).

17. D. O. Kechele et al., Orphan Gpr182 suppresses ERK-mediated intestinal proliferation during regeneration and adenoma formation. J Clin Invest 127, 593–607 (2017).

18. G. Camejo, E. Hurt-Camejo, O. Wiklund, G. Bondjers, Association of apo B lipoproteins with arterial proteoglycans: pathological significance and molecular basis. Atherosclerosis 139, 205–222 (1998).

19. U. Olsson, G. Östergren-Lundén, J. Moses, Glycosaminoglycan-lipoprotein interaction. Glycoconjugate J 18, 789–797 (2001).

20. A. Mehta, M. D. Shapiro, Apolipoproteins in vascular biology and atherosclerotic disease. Nat Rev Cardiol 19, 168–179 (2022).

21. C. D. Schmid et al., GPR182 is a novel marker for sinusoidal endothelial differentiation with distinct GPCR signaling activity in vitro. Biochem Biophys Res Commun 497, 32–38 (2018).

22. S. Melgrati et al., Atlas of the anatomical localization of atypical chemokine receptors in healthy mice. Plos Biol 21, (2023).

23. R. W. Mahley, T. L. Innerarity, S. C. Rall, Jr., K. H. Weisgraber, Plasma lipoproteins: apolipoprotein structure and function. J Lipid Res 25, 1277–1294 (1984).

24. J. P. Scallan, M. Jannaway, Lymphatic Vascular Permeability. Cold Spring Harb Perspect Med 12, (2022).

25. W. O. Dobbins, 3rd, The intestinal mucosal lymphatic in man. A light and electron microscopic study. Gastroenterology 51, 994–1003 (1966).

26. R. J. Torphy, E. J. Yee, R. D. Schulick, Y. Zhu, Atypical chemokine receptors: emerging therapeutic targets in cancer. Trends Pharmacol Sci 43, 1085–1097 (2022).

27. S. Kapas, A. J. Clark, Identification of an orphan receptor gene as a type 1 calcitonin gene-related peptide receptor. Biochem Biophys Res Commun 217, 832–838 (1995).

28. Y. Ikeda, H. Kumagai, A. Skach, M. Sato, M. Yanagisawa, Modulation of circadian glucocorticoid oscillation via adrenal opioid-CXCR7 signaling alters emotional behavior. Cell 155, 1323–1336 (2013).

29. M. Meyrath et al., The atypical chemokine receptor ACKR3/CXCR7 is a broad-spectrum scavenger for opioid peptides. Nat Commun 11, 3033 (2020).

30. C. P. Libeu et al., New insights into the heparan sulfate proteoglycan-binding activity of apolipoprotein E. J Biol Chem 276, 39138–39144 (2001).

31. R. Bonnavion, S. M. Liu, H. Kawase, K. A. Roquid, S. Offermanns, Large chemokine binding spectrum of human and mouse atypical chemokine receptor GPR182 (ACKR5). Front Pharmacol 14, (2023).

32. J. R. Kraehling et al., Genome-wide RNAi screen reveals ALK1 mediates LDL uptake and transcytosis in endothelial cells. Nat Commun 7, 13516 (2016).

33. T. G. Richardson et al., Evaluating the relationship between circulating lipoprotein lipids and apolipoproteins with risk of coronary heart disease: A multivariable Mendelian randomisation analysis. PLoS Med 17, e1003062 (2020).

34. S. Ishibashi et al., Hypercholesterolemia in Low-Density-Lipoprotein Receptor Knockout Mice and Its Reversal by Adenovirus-Mediated Gene Delivery. J Clin Invest 92, 883–893 (1993).

35. M. S. Brown, J. L. Goldstein, A receptor-mediated pathway for cholesterol homeostasis. Science 232, 34–47 (1986).

36. J. B. Weir, New methods for calculating metabolic rate with special reference to protein metabolism. J Physiol 109, 1–9 (1949).

37. J. Folch, M. Lees, G. H. Sloane Stanley, A simple method for the isolation and purification of total lipides from animal tissues. J Biol Chem 226, 497–509 (1957).

38. D. Kraus, Q. Yang, B. B. Kahn, Lipid Extraction from Mouse Feces. Bio Protoc 5, (2015).

39. K. D. Bruce et al., Neuronal Lipoprotein Lipase Deficiency Alters Neuronal Function and Hepatic Metabolism. Metabolites 10, (2020).

40. E. J. Onufer et al., Lipid absorption and overall intestinal lymphatic transport are impaired following partial small bowel resection in mice. Sci Rep 12, 11527 (2022).

