## Supplementary Table and Figures for "GPR182 is a lipoprotein receptor for dietary fat absorption"

**Supplementary Table 1 Mice serum cholesterol, triglycerides and free fatty acid measurements.**

| Gender | Female | | | | | | Male | | | | | |
|---|---|---|---|---|---|---|---|---|---|---|---|---|
| Age (Month) | 2 | | 5 | | 8 | | 2 | | 5 | | 8 | |
| Mouse Genotype | WT (n=10) | GPR182$^{-/-}$ (n=9) | WT (n=10) | GPR182$^{-/-}$ (n=8) | WT (n=10) | GPR182$^{-/-}$ (n=9) | WT (n=10) | GPR182$^{-/-}$ (n=10) | WT (n=10) | GPR182$^{-/-}$ (n=10) | WT (n=8) | GPR182$^{-/-}$ (n=8) |
| Total Cholesterol (mg/dL) | 83.22±7.49 | 97.85±9.08$^{**}$ | 85.28±7.29 | 109.45±7.02$^{****}$ | 93.63±10.15 | 111.72±7.22$^{***}$ | 104.53±5.47 | 121.15±10.30$^{***}$ | 110.17±9.63 | 137.91±17.41$^{***}$ | 115.38±8.53 | 138.64±7.22$^{****}$ |
| Cholesteryl Ester (mg/dL) | 49.93±5.04 | 64.98±6.18$^{****}$ | 51.04±4.11 | 76.55±4.77$^{****}$ | 56.41±7.48 | 77.98±6.28$^{****}$ | 60.39±3.02 | 83.09±9.66$^{****}$ | 63.48±7.15 | 97.64±10.96$^{****}$ | 68.29±5.86 | 97.71±6.94$^{****}$ |
| Free Cholesterol (mg/dL) | 33.29±3.13 | 32.87±3.23$^{ns}$ | 34.24±3.87 | 32.90±5.04$^{ns}$ | 37.22±4.19 | 33.74±3.63$^{ns}$ | 44.14±2.59 | 38.06±3.73$^{***}$ | 46.69±5.68 | 40.27±7.04$^{*}$ | 47.09±5.79 | 40.93±2.94$^{*}$ |
| Triglycerides (mg/dL) | 82.14±5.19 | 75.66±6.37$^{*}$ | 92.39±6.08 | 76.95±5.88$^{***}$ | 104.21±9.37 | 81.54±7.27$^{****}$ | 91.87±6.45 | 79.82±5.07$^{***}$ | 102.85±8.30 | 84.47±6.53$^{****}$ | 119.33±7.78 | 88.15±6.11$^{****}$ |
| Free Fatty Acid (mM) | 0.67±0.05 | 0.61±0.04$^{*}$ | 0.81±0.07 | 0.65±0.05$^{****}$ | 0.91±0.08 | 0.71±0.07$^{****}$ | 0.73±0.05 | 0.69±0.04$^{*}$ | 0.99±0.08 | 0.72±0.06$^{****}$ | 1.18±0.07 | 0.75±0.05$^{****}$ |

### fig. S1

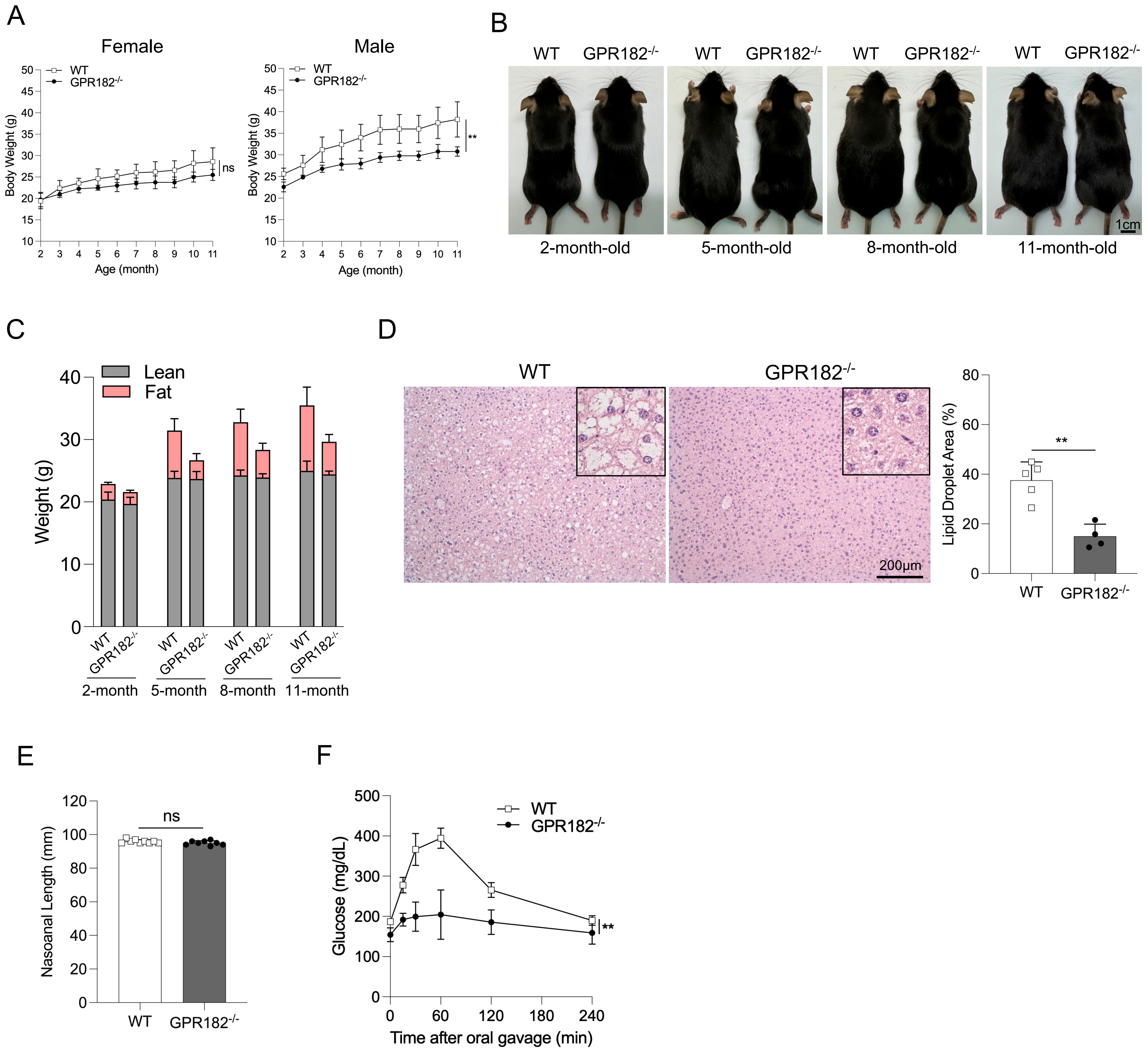


**Supplementary Figure 1 GPR182$^{-/-}$ mice are slimmer and are resistant to diet-induced obesity.**
(A) 2-month-old WT and GPR182$^{-/-}$ mice on regular chow diet were weighed monthly over 9 months. n=5. (B) Representative images of male WT and GPR182$^{-/-}$ mice on chow diet at different ages. (C) Fat and lean masses of male WT and GPR182$^{-/-}$ mice on regular chow diet were assessed by MRI. n=5. (D) H&E staining of livers from 15-month-old male WT and GPR182$^{-/-}$ mice on regular chow diet. Lipid droplet area in livers was quantified. n=4, 5. (E, F) Male WT and GPR182$^{-/-}$ mice were on a 16-week span of HFD, as shown in **Figure 1**. Nasoanal length (E) was determined. n=8, 9. (F) Glucose tolerance in WT and GPR182$^{-/-}$ mice was assessed after 16 weeks of HFD. n=5.

### fig. S2

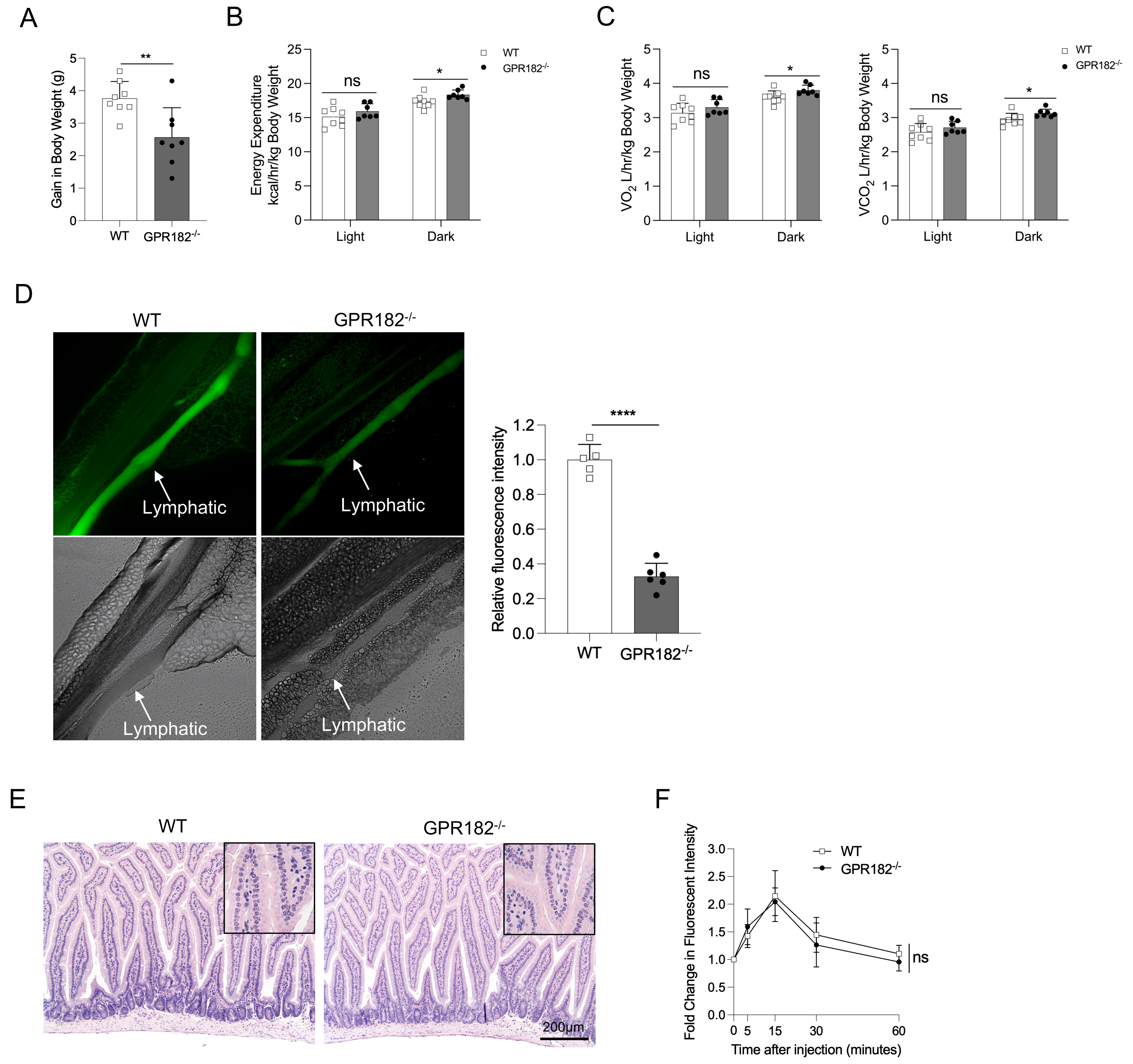


**Supplementary Figure 2 GPR182 is required for dietary lipid absorption.**
(A-C) WT and GPR182$^{-/-}$ male mice at 2 to 3- month- old fed with HFD were assessed for energy balance in two weeks. (A) Body weight gain in a two-week span was quantified. Energy expenditure (B), $O_2$ consumption (C, left) and $CO_2$ production (C, right) were quantified. (D) Representative fluorescent images of mesenteric lymph 2 hours after mice were orally administrated with BODIFY-labeled C16 fatty acid. Fluorescence intensity in mesenteric lymph was quantified. n=5, 6. (E) H&E staining of small intestines from young adult WT and GPR182$^{-/-}$ mice under regular chow diet. (F) Draining function of intestinal lymphatics in adult WT and GPR182$^{-/-}$ mice under regular chow diet was determined by Peyer's patches injection of FITC-Dextran. Right after Dextran injection, blood samples were collected at different timepoints to measure fluorescence. n=5.

### fig. S3

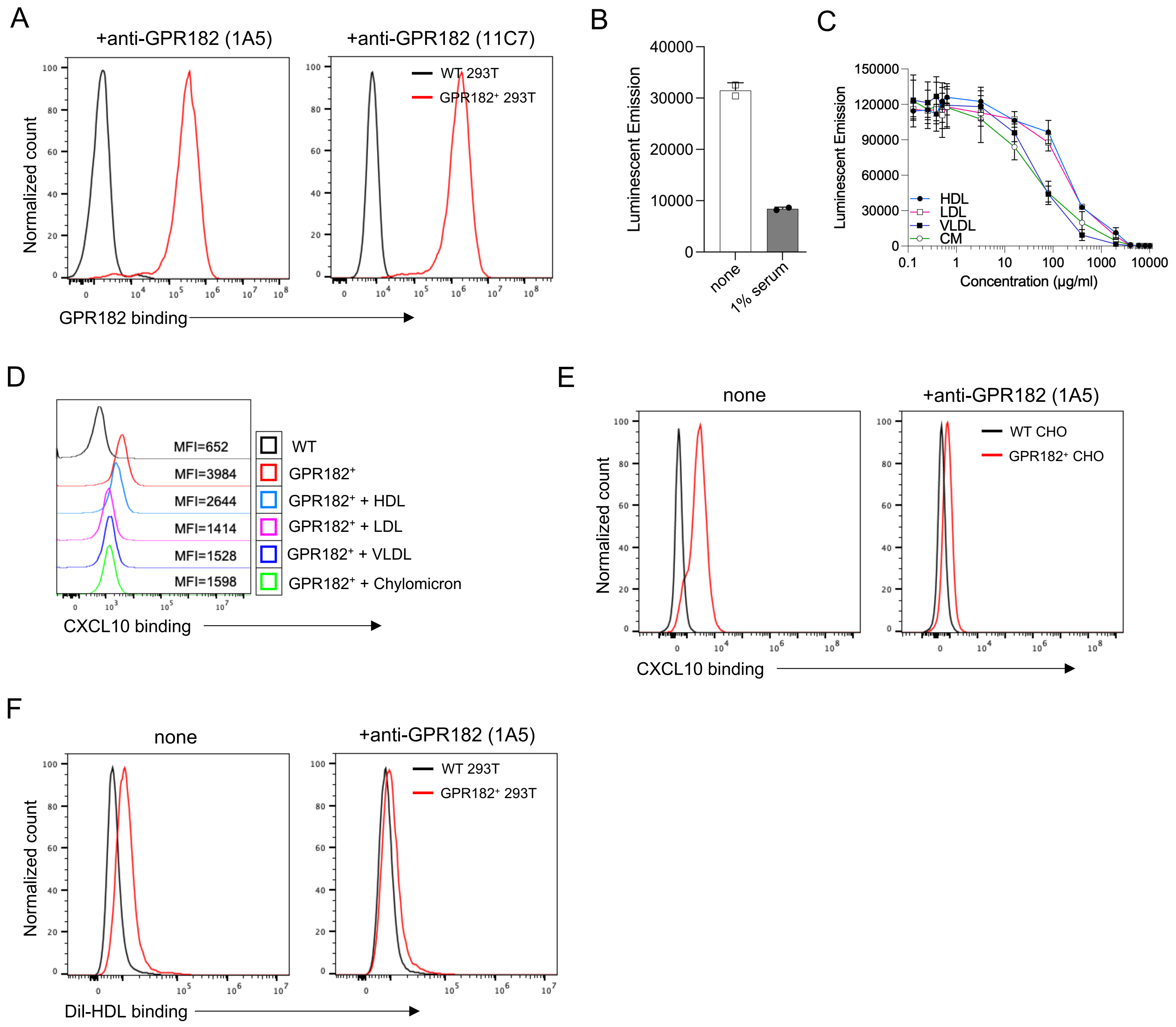


**Supplementary Figure 3 Lipoproteins interact with GPR182.**
(A) The specificity of human GPR182 mAb 1A5 and 11C7 was verified by their specific binding to GPR182-expressing 293T cells. (B) In a NanoBit assay, human serum was added to evaluate its capacity of disrupting the GPR182/β2-arrestin association. (C) Similarly, serum lipoproteins were added at different concentrations to assess their effect in inhibiting the GPR182/β2-arrestin association. (D) Different lipoproteins were tested for their ability to block the binding of $GPR182^{+}$ 293T cells by CXCL10-AF647. (E, F) Human GPR182 mAb (clone 1A5) was assessed for its blocking capacity of GPR182 binding by CXCL10-AF647 (E) or Dil-HDL (F).

### fig. S4

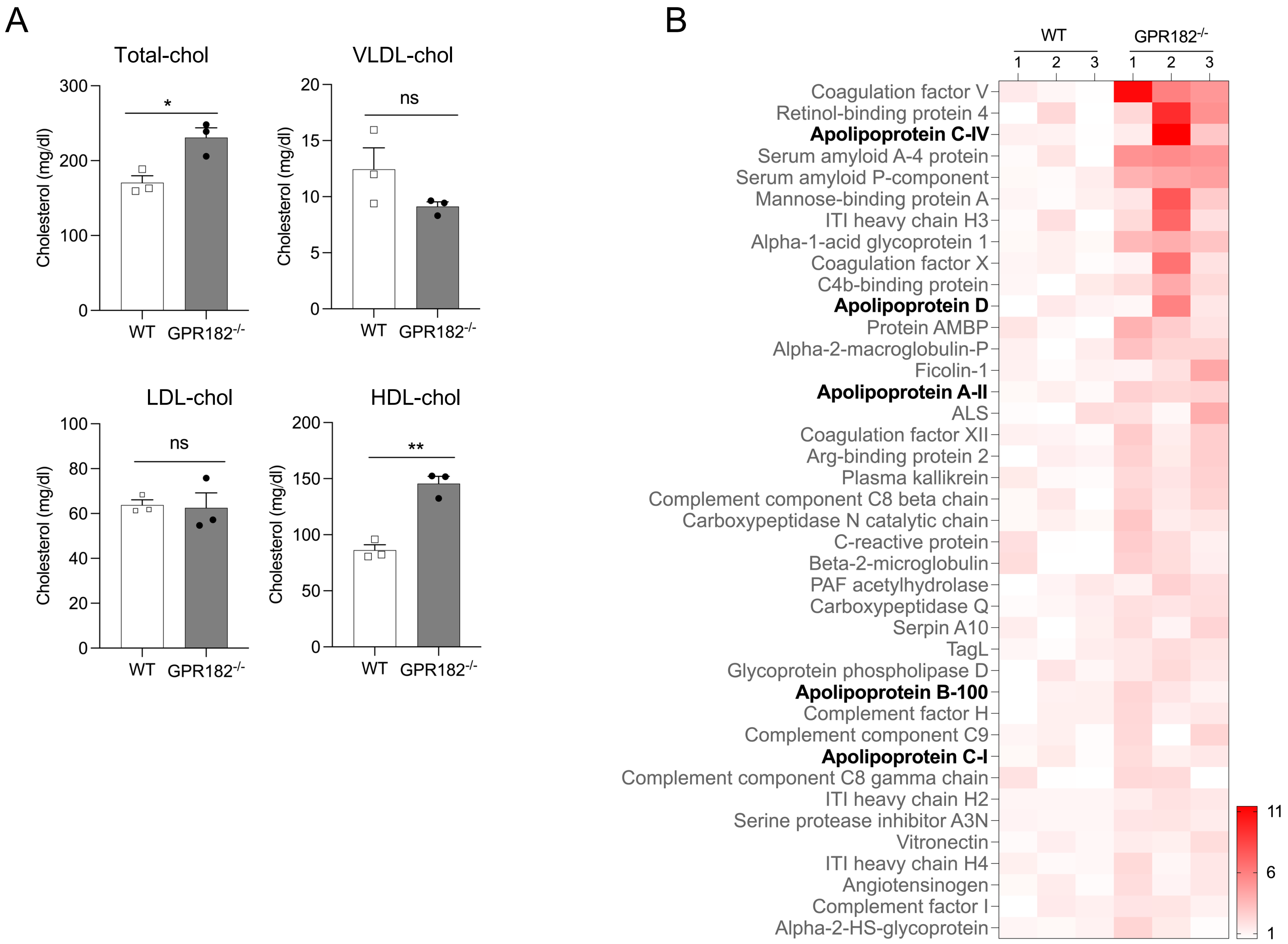


**Supplementary Figure 4 GPR182 regulates lipoprotein homeostasis**. (A) Serum lipoproteins in adult GPR182$^{-/-}$ and control WT mice were quantified by FPLC. (B) Serum proteins in adult WT and GPR182$^{-/-}$ mice were determined by mass spectrometry. The top 40 serum proteins increased in GPR182$^{-/-}$ mice, including several apolipoproteins, were listed. n=3.

#### fig. S5

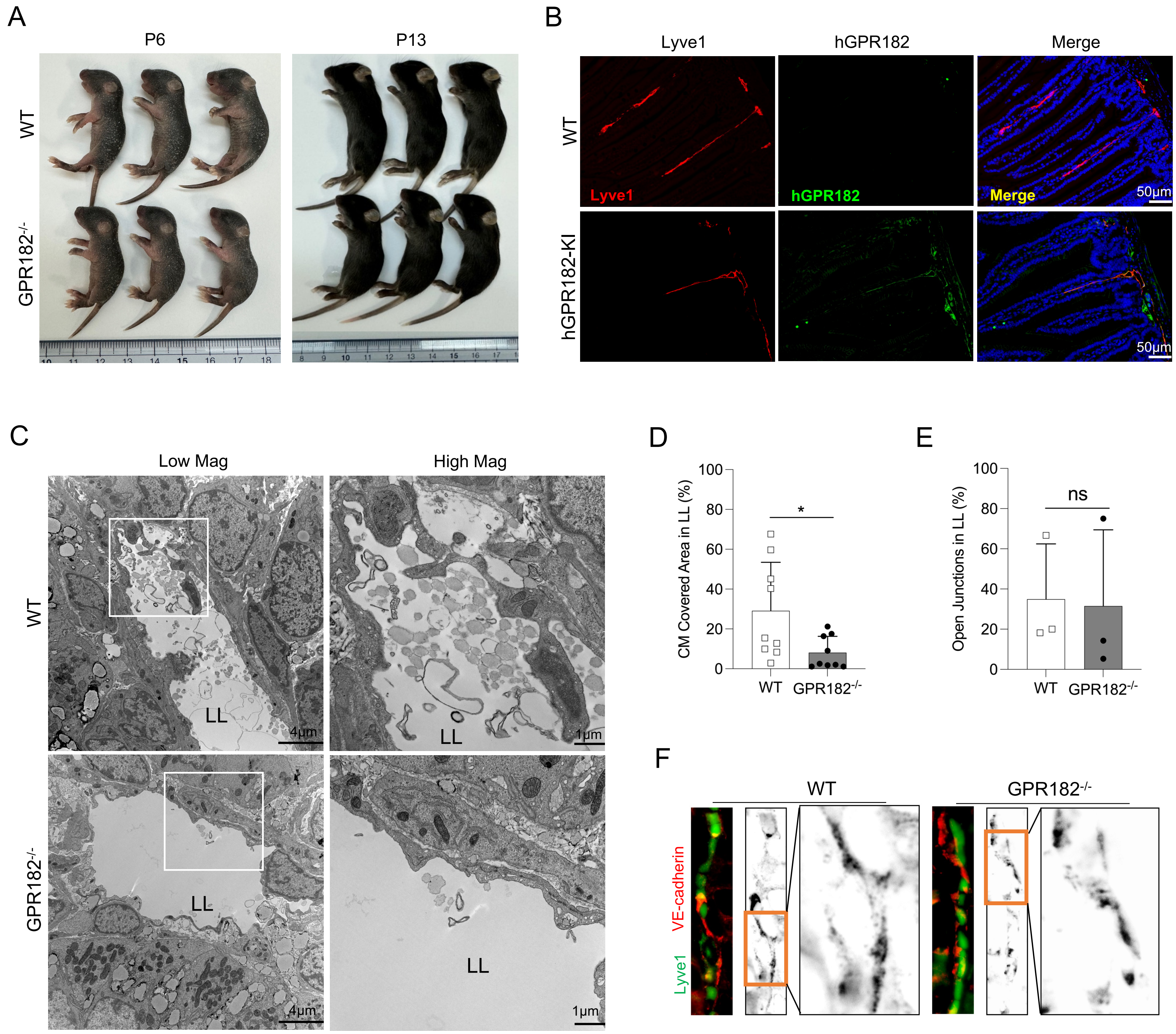


**Supplementary Figure 5 GPR182 on LECs mediates fat absorption in the small intestine.** (A) Images of WT and GPR182$^{-/-}$ newborns (P6, P13) were recorded. (B) Immunofluorescent staining was performed to examine the expressions of hGPR182 and Lyve1 in the small intestine from hGPR182-KI mice. WT mice were used as a negative control for hGPR182 staining. (C) TEM images of intestinal villus from P6 newborns of WT and GPR182$^{-/-}$ mice. LL: lacteal lumen. Quantification of chylomicrons in lacteal lumens (D) and open lacteal junctions (E) was based on TEM images. (F) Representative images of VE-cadherin$^{+}$ LEC junctions of Lyve1$^{+}$ lacteals in the villi of adult WT and GPR182$^{-/-}$ mice.

#### fig. S6

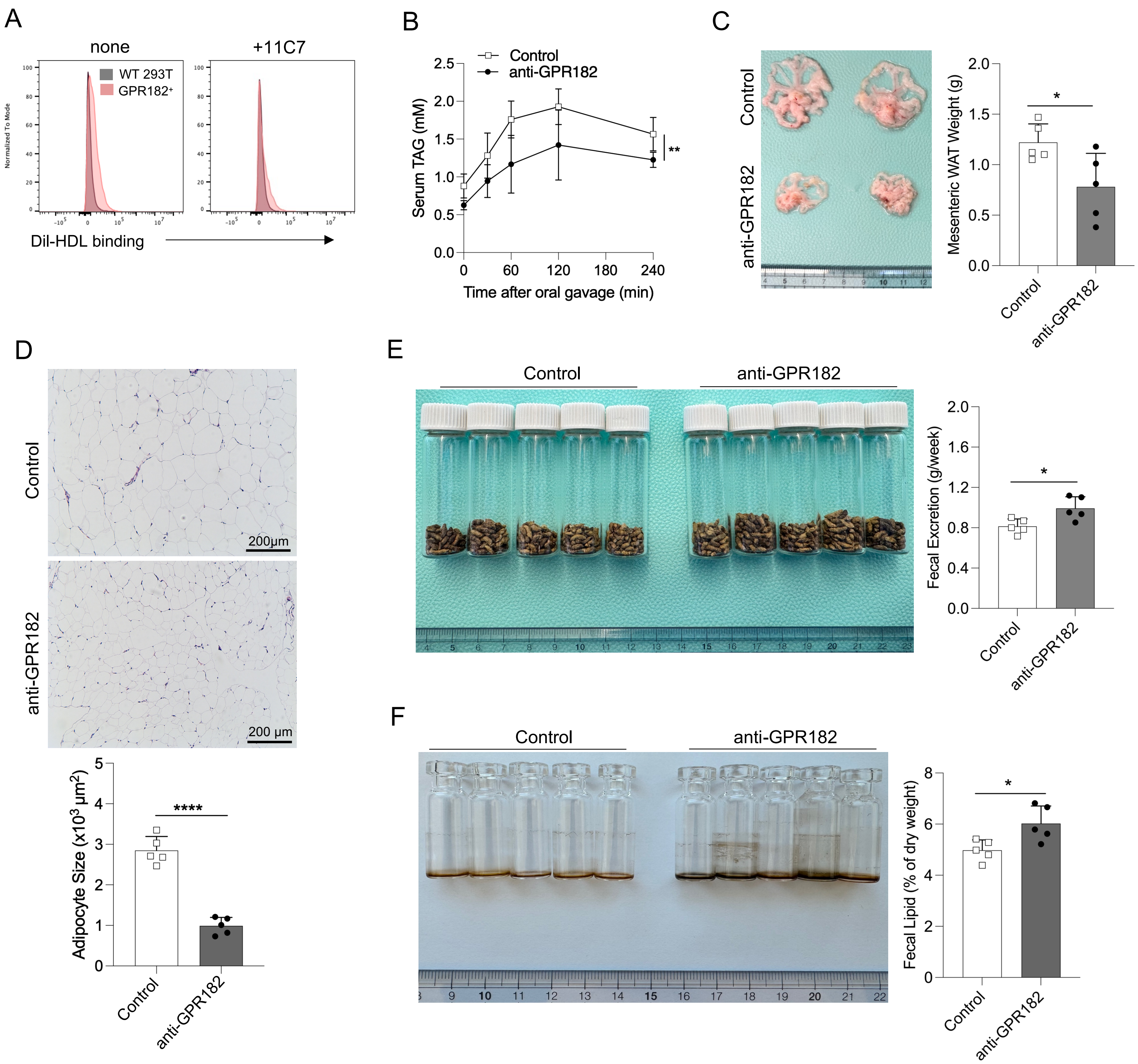


**Supplementary Figure 6 Anti-GPR182 slows down HFD-induced obesity.** (A) Human GPR182 mAb (clone 11C7) was assessed for its capacity of blocking Dil-HDL binding. (B) Human GPR182-knockin (hGPR182-KI) mice pretreated with control or anti-hGPR182 mAb (clone 11C7) were assessed for serum TAG upon olive oil gavage. n=5. (C-F) hGPR182-KI mice in 16 weeks of HFD challenge were treated with control or anti-hGPR182 mAb weekly, as shown in **Figure 5**. n=5. (C) Representative images of mesenteric WATs (mWATs) were shown. mWAT weight was determined in the right panel. (D) H&E staining of mWATs was shown. Adipocyte sizes were quantified. (E) Representative images of feces produced weekly were shown. Weekly fecal excretion was determined in the right panel. (F) Representative images of extracted fecal lipids produced weekly were shown. Fecal lipids were measured in the right panel.

fig. S7

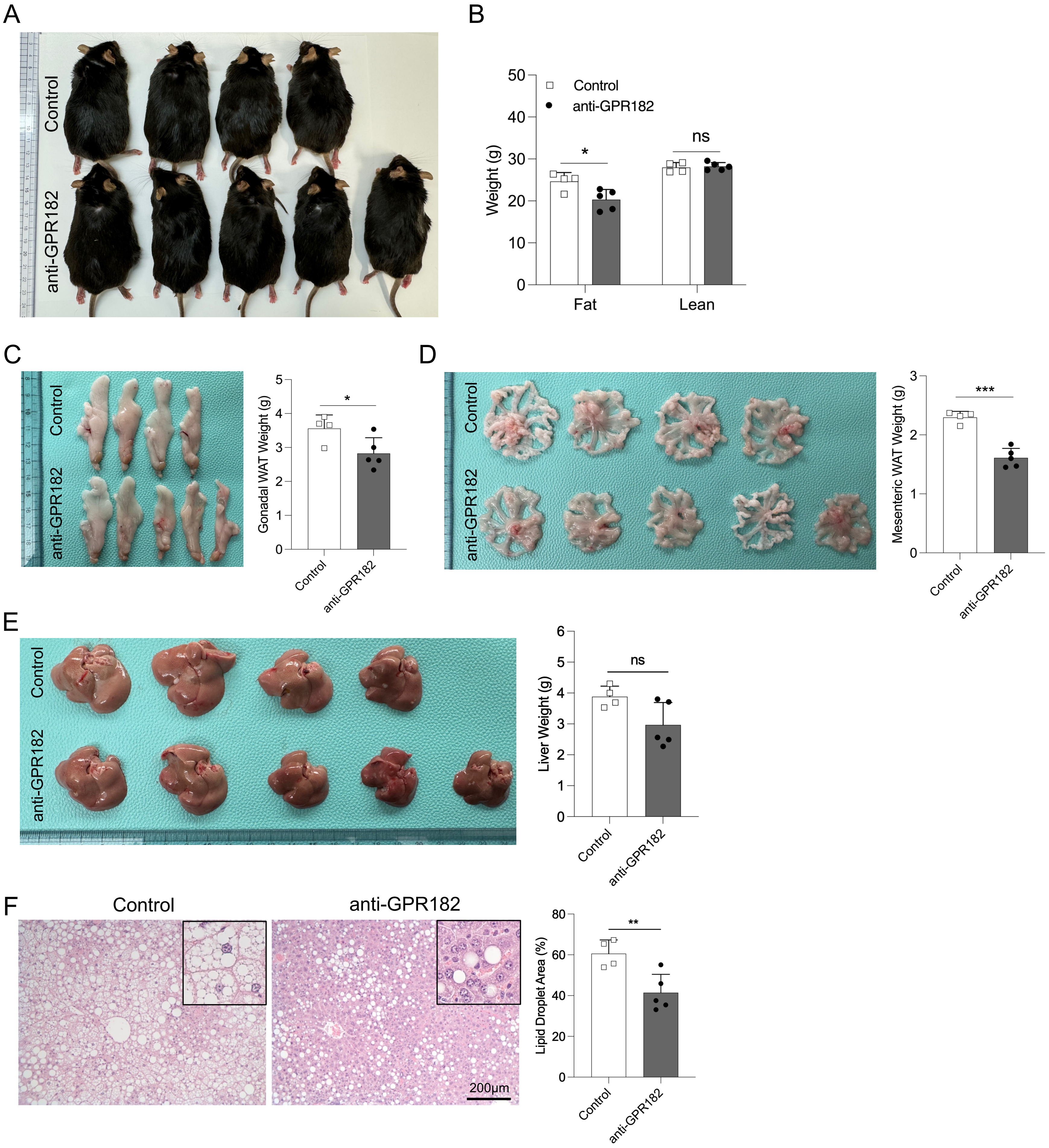


**Supplementary Figure 7 Anti-GPR182 attenuates HFD-induced obesity.** DIO hGPR182-KI mice were treated with control or anti-GPR182 mAb (clone 11C7) twice a week for total 8 weeks. (A) Images of mice after 8 weeks of treatment were shown. (B) Fat and lean weight of mice were determined by MRI. (C) Images of gonadal WATs were shown. Gonadal WAT weight was determined in the right panel. (D) Images of mesenteric WATs were shown. Mesenteric WAT weight was determined in the right panel. (E) Images of livers were shown. Liver weight was determined in the right panel. (F) H&E staining of livers from DIO hGPR182-KI mice with antibody treatment. Lipid droplet area in livers was quantified. n=4, 5.
